# Cultivation in a Natural Microbial Community Enhances the Industrial Performance of a Genetically Engineered Cyanobacterium for Bioplastic Production

**DOI:** 10.1101/2025.08.08.669266

**Authors:** Arianna Zini, Jennifer Müller, Phillipp Fink, Karl Forchhammer

## Abstract

Large-scale production of polyhydroxybutyrate (PHB), a biodegradable bioplastic, using genetically engineered cyanobacteria offers a sustainable alternative to petrochemical-derived plastics. However, monoculture-based phototrophic systems face major limitations, such poor resilience in large-scale reactors, hindering industrial upscaling. To address these challenges, we established a hybrid photosynthetic microbiome by replacing the native cyanobacterium of a natural microbial consortium with a genetically engineered *Synechocystis* strain optimized for PHB production. This new community retained the ecological structure and stability of the original microbiome while gaining synthetic production capacity. Compared to the axenic strain, the hybrid system exhibited enhanced robustness under abiotic stress, including light and temperature fluctuations, and improved tolerance to operational instability. These features made it suitable for upscaling and application in non-sterile environments. The hybrid microbiome sustained PHB production in scaled photobioreactors, reaching up to 32% PHB per cell dry weight (CDW) equal to ∼230 mg L^-1^ under fully photoautotrophic conditions. Production was also achieved under dark conditions with acetate supplementation, highlighting the system’s metabolic flexibility. This work demonstrates the successful integration of an engineered phototroph into a stable native microbiome, positioning hybrid communities as powerful platform for industrial biotechnology.

## Introduction

Polyhydroxybutyrate (PHB), a member of the polyhydroxyalkanoate (PHA) family, is a biodegradable polymer synthesized by various bacteria and archaea as an intracellular carbon and energy store under nutrient-limited conditions (Gundlapalli and Ganesan, 2025; McAdam et al., 2020). Its favourable mechanical properties, combined with its biodegradability in biologically active environments, make PHB a promising substitute for petrochemical-based plastics (Gautam et al., 2024; McAdam et al., 2020).

*Synechocystis* sp. PCC 6803 (*Synechocystis* from now on) is a well-established model cyanobacterium and a natural PHB producer. Under nitrogen limitation, *Synechocystis* undergoes chlorosis, a dormant state characterized by degradation of the photosynthetic apparatus and accumulation of storage carbohydrates, namely glycogen and PHB (Doello et al., 2021; Forchhammer and Schwarz, 2019). Leveraging this, in previous work, we characterized and engineered the metabolic pathways sustaining PHB synthesis (Orthwein et al., 2021). This resulted in generation of a new strain termed *Synechocystis* sp. PCC 6803 PPT1 (PPT1 from now on), capable of accumulating up to 60% PHB per cell dry weight (CDW) during nitrogen-starvation when NaHCO_3_ was used as a carbon source, and 80% when acetate was used (Koch et al., 2020). However, due to the low biomass in the cultures, overall productivity was limited to 5 mg L⁻¹ d⁻¹. A key limitation to the scalability of cyanobacteria are the high production costs, driven by low product yields, expensive equipment and significant maintenance requirements (Price et al., 2022; Rueda et al., 2023). Moreover, in large, high-density cultures, limited light penetration hampers the onset of chlorosis, making PHB accumulation difficult even after complete nitrate consumption (Klotz et al., 2015). Monocultures are sensitive to environmental stresses common in industrial-scale cultivation, such as fluctuating light intensity, temperature shifts, mechanical disturbances and occasional invasion by foreign microorganisms that outcompete cyanobacteria (Awasthi et al., 2014; Huang et al., 2017). These challenges add on top of the primary limitation associated with cyanobacterial scalability: high production costs.

Diverse microbial communities are generally more resilient to environmental stress and support broader metabolic functionality than monocultures (Awasthi et al., 2014; Jousset et al., 2011). Therefore, synthetic and engineered consortia are emerging as valuable tools in industrial biotechnology, offering improved adaptability and productivity (Giri et al., 2020). Cyanobacteria-enriched microbiomes derived from natural environments have demonstrated robustness under non-sterile conditions in pre-industrial photobioreactors, making them a cost-effective platform for large-scale applications (Altamira-Algarra et al., 2024).

We hypothesized that replacing the native phototroph in a stable, natural microbiome with a genetically engineered cyanobacterium could overcome the limitations of axenic cultures. To test this, we introduced the strain PPT1 into a protective microbial community, creating what we term a *hybrid microbiome*: a native microbial consortium in which certain species have been selectively substituted with engineered ones. This strategy preserves the ecological integrity of the original community while enabling new synthetic functions, offering a robust platform for advancing phototrophic biotechnologies.

## Experimental Procedures

### Cyanobacterial cultivation conditions

All cyanobacterial cultures, axenic and community-based, were pre-cultured in BG11 medium supplemented with 5 mM NaHCO₃ under continuous shaking (120 rpm) at 28 °C and light intensity of ∼50–60 µmol m⁻² s⁻¹ (Rippka et al., 1979). Antibiotics (kanamycin and spectinomycin, 50 µg mL⁻¹ each, depending on the strain’s resistance) were included for axenic cultivation of mutant strains but omitted in microbiome cultures. Pre-cultures at OD₇₅₀ ∼1 were harvested by centrifugation (15 min, 4200 rpm) and resuspended in BG11_0_ medium (BG11 lacking NaNO₃). The traditional two-stage PHB production process, comprising an initial growth phase in BG11 followed by re-inoculation of the pellet into BG11_0_, was replaced with a simplified one-step protocol. Cultures were directly inoculated into BG11_0_ supplemented with 4-8 mM NaNO₃ and NaHCO₃ as a carbon source at a starting OD₇₅₀ of 0.2. As nitrate became limiting, cells transitioned into chlorosis and initiated PHB biosynthesis, thus allowing the need for medium exchange to be circumvented.

Small-scale experiments were conducted in 100 mL Erlenmeyer flasks containing 50 mL of culture. For large-scale setups, either 12 L borosilicate glass vessels (cultivation volume: 8–10 L) or custom flat-panel reactors (dimensions: ∼50 × 26 × 3 cm; volume: 4 L) were used. Sterile conditions involved standard aseptic techniques, including the use of autoclaved media and laminar flow handling. Non-sterile conditions were mimicked by inoculating cultures in ambient air using non-autoclaved BG11 medium. For cultivation on solid medium, cultures were grown on BG11 Difco-agar 1.5% supplemented with 5 mM NaHCO₃ to assess colony formation, or on BG11 supplemented with glucose and casamino acids for contamination screening.

A list of the strains and microbial communities used in this study is provided in the Supporting Information in Table 1.

### Selective isolation of heterotrophic community members

The Wild Cyanobacteria-Rich Microbiome (WCRM) we used, previously described (Altamira-Algarra et al., 2024), is characterized by the presence of diverse cyanobacterial taxa, most of which belonged to the *Synechocystis* sp. To enrich and isolate the heterotrophic fraction of the WCRM, cultures were plated on BG11 prepared with agar-agar (Bernd Euler Biotechnologie-Mikrobiologie), instead of Difco-agar. Plates were incubated at 28 °C under dim light (10–20 µmol m⁻² s⁻¹) for 2–4 days (Fig. 1a). This approach enabled the selective proliferation of the non-cyanobacterial part of the community. Growth was supported by residual organic compounds from the inoculum, derived from cyanobacterial exudates during prior liquid cultivation. For qualitative assessment of heterotrophic diversity and abundance, culture aliquots were also plated on LB agar (Luria Bertani, Lennox).

**Figure 1:**
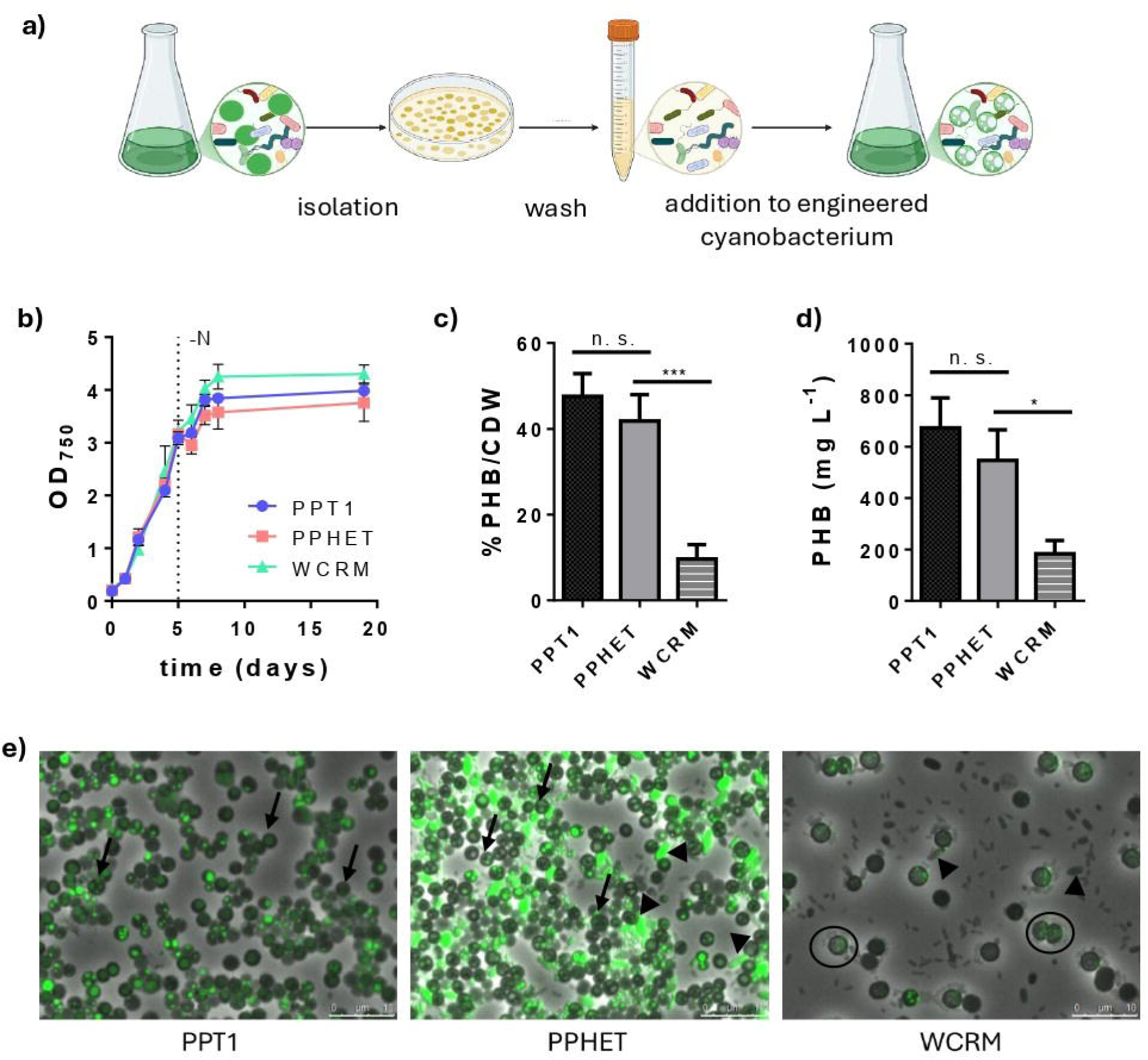
Generation and growth of hybrid cyanobacterial microbiomes. **a)** Schematic representation of the method used to generate hybrid microbiomes by replacing the native cyanobacterial component of the wild-type community with a genetically engineered strain. **b)** Growth curve of PPT1 (blue), PPHET (pink) and a wild cyanobacteria-rich microbiome (WCRM, green) over 19 days in a one-step cultivation PHB production process. The dotted line represents the point at which nitrate (NO_3_ ^-^) had been totally consumed; **c-d)** PHB production of PPT1, PPHET and WCRM at the end of the process in terms of %PHB/CDW (**c**) and in mg L^-1^ (**d**); data points and bar columns represent the mean of three biological replicates at each time point ±SD. For the PHB evaluation, statistical significance was determined using one-way ANOVA with Brown-Forsythe test for variance and Dunnett’s multiple comparisons test against the control group (PPHET). **e)** Microscopic pictures of PPT1, PPHET and WCRM at the end of the process, following staining of PHB granules with BODIPY (green). Images show overlays of phase-contrast and GFP fluorescence channels. Arrows indicate PPT1 cells with PHB stained granules in the axenic (PPT1) and the hybrid microbiome (PPHET) culture. Arrowheads indicate non-*Synechocystis* members of the community in PPHET and WRCM which display stained granules. Circles indicate the wild *Synechocystis* originally present in the natural community.

### Microscopy and staining procedure

Fluorescence microscopy was performed on a LeicaDM5500B (Leica Microsystems, Germany) with objective lenses and filter cubes. PHB granules were visualized via BODIPY staining using the GFP channel, with a modified method previously described (Fink et al., 2025). Briefly, 1.5 µL of BODIPY dye solution (1 mg mL^-1^ in DMSO) was added to 100 µL of cyanobacterial culture. After 10 minutes of incubation in the dark at room temperature, cells were washed two times with phosphate-buffered saline (PBS) and then applied on microscope slides pre-coated with 1% (w/v) agarose to immobilize cells for imaging.

### Chlorophyll *a* quantification

Chlorophyll *a* (Chl *a*) content was quantified using methanol extraction. A 1 mL culture sample was centrifuged at 14,000 rpm for 5 minutes, and the supernatant was discarded. The pellet was resuspended in 1 mL of 90% methanol and incubated in the dark at 4 °C for 30 minutes to allow pigment extraction. After incubation, samples were centrifuged again at 14,000 rpm for 5 minutes, and the absorbance (A) of the supernatant was measured at 665 nm using 90% methanol as a blank. Chlorophyll *a* concentration was calculated using the following equation: Chl *a* (µg/mL) = A₆₆₅ × 13.9.

### Nitrate (NO_3_ ^-^ ) determination

Nitrate concentrations were qualitatively monitored using semi-quantitative test strips (QUANTOFIX® Nitrate/Nitrite, Macherey-Nagel GmbH & Co. KG, Düren, Germany). These strips provide a detection range of 10–500 mg L^-1^ for NO₃⁻ and 1–80 mg L^-1^ for NO₂⁻. Measurements were used to approximately estimate nitrogen depletion during cultivation.

### pH measurements

The pH of culture samples was measured using a Mettler Toledo FiveEasy™ pH/mV meter equipped with an LE422 microelectrode. The electrode has a measurement range of pH 0–14 and is suitable for temperatures up to 80 °C.

### PHB quantification

The method utilized was previously described (Fink et al., 2025). Shortly, 5-10 mL bacterial culture was harvested, cell dry weight was determined and subsequently boiled in 1 ml of concentrated H_2_SO_4_ (18 M) for 1 h at 100 °C to convert PHB to crotonic acid. The boiled cell culture was diluted 1:20 with 0.014 M H_2_SO_4_ and centrifuged at 20,000 g for 5 min at RT. The supernatant was analyzed by HPLC (HITACHI Chromaster, VWR, Germany) using a Nucleosil 100-5 C18 reverse-phase column (125 × 3 mm, 5 μm, 100 Å) fitted with a precolumn (4 x 3 mm). Samples (5 μL) were eluted isocratically at 1 mL/min using 30:70 MeOH/20 mM phosphate buffer (pH 2.5) for 10 min, and crotonic acid was detected at 210 nm. PHB standards and crotonic acid calibration curves were used for quantification via linear regression.

### Statistical Analysis

Small-scale experiments were performed in two or three biological replicates, as indicated in the corresponding figures or sections. Experiments conducted in duplicate at small scale were qualitatively validated through repetition in upscaled systems to assess consistency in system performance. Data are presented as mean values, with error bars representing the standard deviation (±SD). A *p*-value of less than 0.05 (*p* < 0.05) was considered statistically significant. Statistical analyses were performed using GraphPad Prism version 6. Depending on the experimental design, either one-way or two-way ANOVA was applied. For one-way ANOVA, the Brown-Forsythe test was used to assess equality of variances, followed by Dunnett’s multiple comparisons test to compare experimental groups to a control. For two-way ANOVA, Sidak’s multiple comparisons test was used to evaluate differences between groups.

### 16s library preparation and sequence analysis

Samples were collected from the respective cultures 1 week after inoculation in full BG11. Sample preparation and sequencing was done at Core Facility Genomics/NCCT Microbiology, Institute for Medical Microbiology and Hygiene, University Hospital Tübingen. Genomic DNA was isolated using the Quick DNA Fungal/Bacterial Miniprep Kit (Zymo Research) according to the manufacturer’s protocol, with 10 min. bead beating in ZR BashingBead Lysis Tube (Zymo Research). The resulting DNA was quantified with a Qubit dsDNA BR/HS Assay Kit (Thermo Fisher). Libraries for 16S amplicon sequencing were prepared in a two-step PCR approach: using KAPA HiFi HotStart ReadyMix (Roche), primers 515F and 806R (Caporaso et al., 2011) (∼350 bp fragment of the 16S V4 region), and index primer mix (IDT for Illumina DNA/RNA UD Indexes, Tagmentation). The pool was sequenced on an Illumina MiSeq device using the MiSeq Reagent Nano Kit v2 (500 cycles) with 2 x 250 bp read length and a depth >100k reads per sample.

Date were demultiplexed with the nf-core/demultiplex v1.4.1 pipeline (Ewels et al., 2020) and further processed with the nf-core/ampliseq v2.11.0 pipeline (Straub et al., 2020) of the nf-core collection of workflows (Ewels et al., 2022). Both pipelines use reproducible software enviroments from Bioconda (Grüning et al., 2018) and Biocontainers projects (Da Veiga Leprevost et al., 2017).

Data quality was evaluated, and all primer sequences were trimmed. Adapter and primer-free sequences were processed sample-wise (independent) with DADA2 v1.30.0 (Callahan et al., 2016). Taxonomic classification was performed by DADA2 and the database ‘Silva 138.1 prokaryotic SSU’ (Quast et al., 2012). ASV (Amplicon Sequence Variants) sequences, abundance and DADA2 taxonomic assignments were loaded into QIIME2 v2023.7.0 (Bolyen et al., 2019).

## Results

### Generation of the hybrid microbiome PPHET with the cyanobacterium PPT1 as keystone species

To construct a stable microbiome, we combined the genetically engineered cyanobacterial strain *Synechocystis* sp. PCC 6803 PPT1 (hereafter PPT1) with the non-cyanobacterial fraction of a wild cyanobacteria-rich microbiome (WCRM), previously described in Altamira-Algarra *et al*., 2024. After the isolation of the non-cyanobacterial fraction from WCRM (heterotrophs, hereafter), colonies were resuspended in BG11, serially diluted (*het A* to *het D*), and used to inoculate exponentially growing axenic PPT1 cultures (Fig. 1a; Supporting information, Fig. S1a). We refer to the new hybrid culture with PPT1 as core species as PPHET consortia. As a control, *Synechocystis* sp. PCC 6803 expressing eGFP (referred to as GFP) (Orthwein et al., 2021) was used in a parallel setup to assess the effect of replacing native cyanobacteria and to validate the engineering approach. No significant differences were observed between growth in communities and in isolation for both PPT1 and GFP (Supporting information, Fig. S1a), indicating that the hybrid system did not impair cyanobacterial growth. Chlorophyll *a* content was also quantified and found to be comparable across all conditions, further confirming active phototrophic growth (Supporting information, Fig. S1b). Microscopy analysis validated the successful colonization and replacement of the original phototrophic strain with PPT1 or GFP (Supporting information, Fig. S1c). Despite variations in the density of the initial heterotrophic inoculum, the resulting communities converged toward similar structural compositions over time, as evidenced by consistent colonies morphology and abundance on LB agar plates (Supporting information, Fig. S1d), suggesting that the metabolic interactions with the cyanobacterium shape the community composition. To confirm the stable integration of PPT1 within the hybrid microbiome, we performed multiple assessments, including culture plating on selective media, repeated propagation cycles, and colony PCR. These results are presented in the Supporting information (Supporting information, Fig. S1d-e).

To evaluate growth dynamics and PHB production, we compared PPT1, PPHET and WCRM. In this experiment, cultures were inoculated in BG11_0_ medium supplemented with 5 mM of NaNO_3_ and 10 mM of NaHCO_3_. Upon nitrogen depletion, 50 mM of NaHCO_3_ were added, to boost PHB accumulation. The experiment was conducted in three biological replicates over 19 days, comprising the initial growth phase followed by chlorosis and PHB accumulation. The final biomass accumulation was comparable in all three systems, with OD₇₅₀ values reaching approximately 4 (Fig. 1b). PHB accumulation was assessed both quantitatively and qualitatively. PPT1 accumulated 50% PHB per cell dry weight (CDW). PPHET reached similar PHB levels, up to 48% PHB/CDW, whereas the WCRM system, comprising wild-type *Synechocystis*, produced around 10% PHB/CDW (Fig. 1c). PHB titers and productivity were also evaluated, with PPT1 and PPHET achieving similar performances exceeding 600 mg L^-1^ (Fig. 1d), corresponding to a productivity of approximately 35 mg L⁻¹ d⁻¹.

Fluorescence microscopy with BODIPY staining enabled visualization of intracellular PHB granules (Fig. 1e). Comparable PHB accumulation was observed in PPT1 under axenic and community conditions (indicated by black arrows). Interestingly, in PPHET not only PPT1 but also other members of the community stained positive for BODIPY, suggesting they may be able to accumulate PHB as well (indicated by black arrowheads in PPHET and WCRM). As expected, wild-type *Synechocystis* in WCRM accumulated fewer and smaller PHB granules (indicated by a black circle). It is also worth noting that non-*Synechocystis* members of the community in PPHET accumulated more PHB than to WRCM, suggesting that different interactions may occur in the wild microbiome compared to the hybrid one. Additionally, microscopy analysis revealed that wild-type *Synechocystis* cells in WCRM are larger than the engineered PPT1 cells, offering a visual mean to distinguish the engineered strain from the wild-type within mixed communities.

### Microbial dynamics and ecological insights from 16S rRNA sequencing

To assess microbial diversity in PPHET compared to WCRM microbiomes, samples were collected one week after inoculation in BG11 and standard growing conditions for 16S rRNA amplicon sequencing. The evaluation of data quality revealed that 93.3% of the sequences per sample passed the filtering. After denoising and removal of chimeric reads between 70.88% and 87.19% reads per sample (average 82.5%) were used to determine the amplicon sequencing variants (ASVs). 139 amplicon sequencing variants (ASVs) were obtained across all samples.

Overall, the two communities shared 80–90% of their taxonomic composition. At the phylum level, both were dominated by Cyanobacteria, Proteobacteria, and Bacteroidota, with Cyanobacteria representing ∼70% of the total community (Fig. 2a). The predominant classes included Cyanobacteriia, Alphaproteobacteria, and Bacteroidia. Using the revised cyanobacterial taxonomy according to whole-genome phylogenomic analyses (Salazar et al., 2020), we performed order-level taxonomy analysis, which revealed key differences between the two communities. While both PPHET and WCRM contained Cyanobacteriales, reflecting the presence of *Synechocystis*, WCRM also included additional cyanobacterial orders, such as the unicellular Thermosynechococcales and in minor abundancy the filamentous Phormidesmiales and Leptolyngbyales (Supporting Information, Table 2), suggesting a multi-phototrophic community. In contrast, PPHET contained only Cyanobacteriales, consistent with the introduction of a single engineered strain (PPT1) (Fig. 2b). Microscopic observations further supported this finding, confirming PPT1 as the dominant, and likely sole, cyanobacterial species in PPHET (blue arrowheads, Fig. 2c; black arrows, Fig. 1e). These analyses confirm the effectiveness of our approach to selectively retain heterotrophs during community assembly. Despite the overall compositional similarity, ∼10% of the community structure differed between WCRM and PPHET. WCRM exhibited higher abundancy in Pirellulales, Acetobacterales, Phycisphaerales, Deinococcales, and Burkholderiales, whereas PPHET was more abundant in Chitinophagales, Cytophagales, and Sphingobacteriales (Fig. 2b). Groups of microorganisms with a relative abundancy lower than 0,05% were grouped in “Others” and can be found in the Supporting Information, Table 2. These shifts likely reflect selective pressures imposed by the presence of the engineered cyanobacterium.

**Figure 2:**
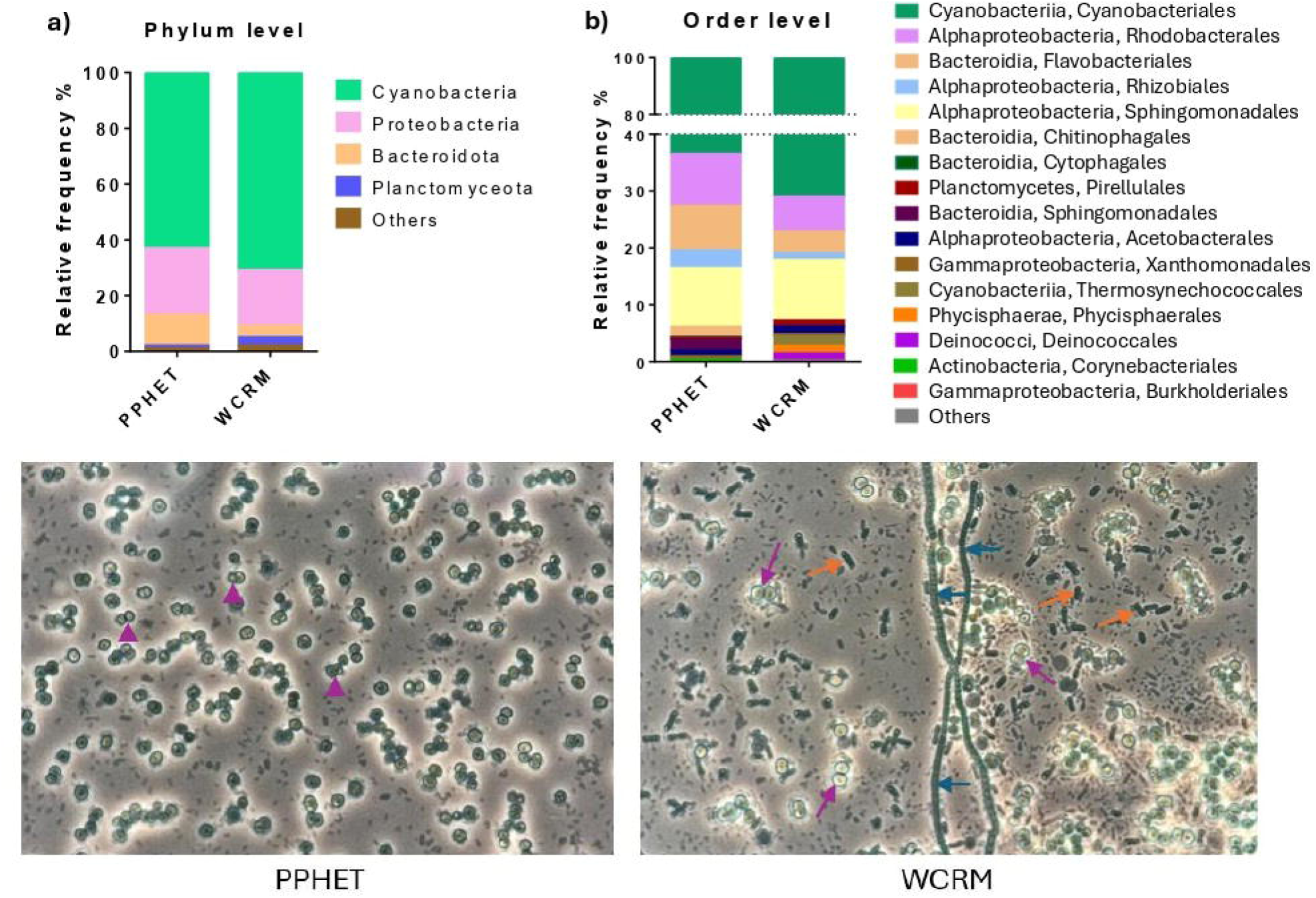
Microbial Community Composition of PPHET and WRCM 7 days after inoculation in BG11. **a-b)** Comparison of the biodiversity composition in PPHET and WCRM at the phylum level **(a)** and at the order level **(b)**. Stacked bar plots show the relative abundance of taxonomic groups within each community. Each bar represents one sample, with segments indicating the proportion of each group. **c)** Microscopic pictures of PPHET and WCRM during normal growth in standard BG11. Samples were imaged using transmitted light brightfield microscopy and acquired via eyepiece projection. Purple arrowheads indicate *Synechocystis* PPT1, which appears smaller than *Synechocystis* from the WCRM (purple arrows). In WCRM, orange arrows indicate members of the Thermosynechococcales order, whereas blue arrows indicate filamentous cyanobacteria, likely members of the Phormidesmiales or of the Leptolyngobyales order, according to our 16S rRNA results.

### Robustness of the hybrid microbiome under biotic and abiotic stress factors

To evaluate the robustness of PPHET compared to the axenic strain PPT1, both systems were tested under various biotic and abiotic stress conditions, including non-sterile environments, elevated light intensities and temperature fluctuations. PPT1 and PPHET were cultivated in BG11_0_ medium supplemented with 4 mM NaNO₃ and 10 mM NaHCO₃, without any carbon boost upon nitrogen depletion (Supporting information, Fig. S2). Under solely non-sterile conditions, both cultures exhibited moderate reductions in PHB content, but PPT1 maintained growth and PHB accumulation, suggesting that microbial contamination played a minor role in small-scale systems.

To evaluate the effect that abiotic factors have on culture fitness, the experiments were conducted under sterile conditions. To determine the effect of high light stress, light intensity was gradually increased to 400 µmol m⁻² s⁻¹. Under these conditions, both systems exhibited similar pH trends (Fig. 3b), but their physiological responses diverged. PPT1 growth was strongly inhibited under high light, with OD₇₅₀ decreasing to ∼2 by day 21 and PHB production limited to ∼100 mg L^-1^ (Fig. 3a, 3c). In contrast, PPHET maintained pigmentation and stable biomass (OD₇₅₀ ∼5) with PHB titers averaging ∼300 mg L^-1^, indicating improved stress tolerance and continued carbon flux toward PHB synthesis (Fig. 3c). These results suggest that heterotrophic members of PPHET may alleviate oxidative stress by consuming excess oxygen.

**Figure 3:**
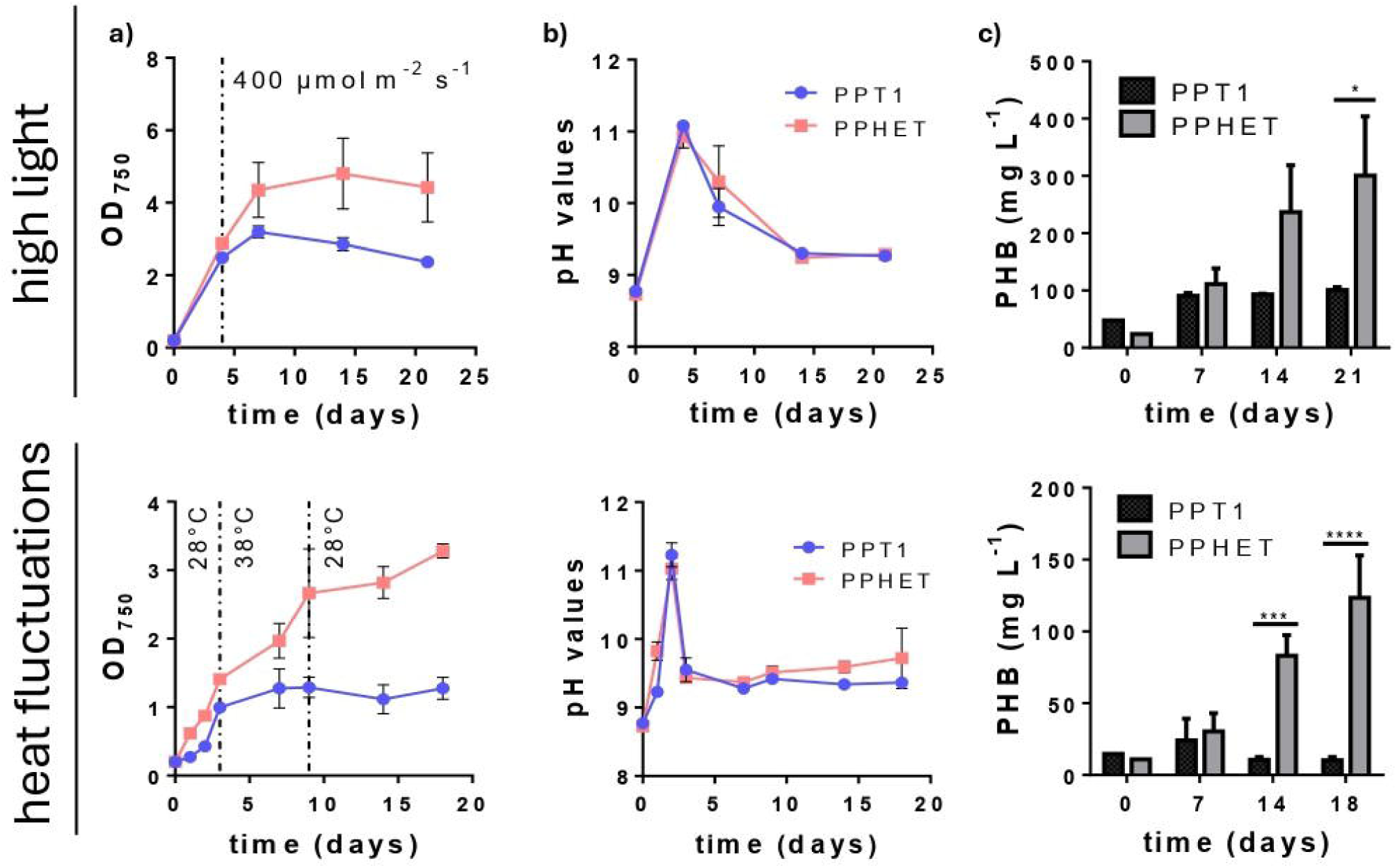
Improved robustness of PPHET compared to PPT1. Comparison of PPHET (pink) and PPT1 (blue) under high light (top) and heat fluctuations (bottom). **a)** Growth curve. Dashed lines indicate the point at which 400 µmol m⁻² s⁻¹ was reached (top) and when the temperature was shifted (bottom). **b)** pH behavior overtime. **c)** PHB production evaluated in mg L^-1^. Data points and column bars represent the mean of at least two biological replicates at each time point ±SD. For PHB data analysis, statistical significance was determined using two-way ANOVA followed by Sidak’s multiple comparisons test. Only statistically significant differences are indicated. These observations are further verified in big scale in the next section.

To assess thermal tolerance, cultures were exposed to a transient heat shock (38 °C) followed by a return to optimal conditions. PPT1 exhibited irreversible bleaching and growth arrest, with no recovery upon temperature normalization (Fig. 3a; Supporting information, Fig. S3). In contrast, PPHET maintained pigmentation during heat stress and resumed growth upon return to standard conditions. PHB production remained detectable in PPHET (up to 150 mg L^-1^), whereas it was negligible in PPT1 (Fig. 3c). These results indicate that PPHET confers resilience to transient heat stress and supports functional recovery.

### Metabolic flexibility of the hybrid microbiome

Upscaling PHB production in axenic cyanobacterial cultures is limited not only by poor robustness but also by the requirement for chlorosis as a trigger for PHB accumulation. Chlorosis is induced by nitrogen depletion combined with the presence of sufficient light (Klotz et al., 2015), a condition that is difficult to achieve uniformly at scale, particularly in large reactors where the surface-to-volume ratio limits effective illumination. Without it, cultures fail to enter chlorosis and do not accumulate PHB.

According to our microscopic results, the two microbiomes not only rely on their cyanobacteria population for PHB production, but also on other members of the community (Fig. 1e). Assuming that the PPHET community is composed exclusively of *Synechocystis* PPT1, we concluded that the other members of the community that are stained with BODIPY in Fig. 1e (indicated by black arrowheads), are non-cyanobacterial species, likely capable of a heterotrophic metabolism. Supporting this view, Altamira-Algarra and colleagues (Altamira-Algarra et al., 2024), showed that in the WCRM, PHB production can be induced by first allowing photoautotrophic growth and then providing sodium acetate under dark incubation conditions. We reasoned that a similar approach might also work in PPHET.

To test this, we applied a heterotrophic induction strategy to PPHET. Following photoautotrophic growth, cultures were transferred to BG11_0_ supplemented with 30 mM acetate and 10 mM NaHCO₃ and incubated under either continuous light (as a control) or complete darkness. Under light, PPHET achieved ∼55% PHB/CDW compared to ∼45% in PPT1. In darkness, only PPHET continued to accumulate PHB, reaching ∼38% PHB/CDW, whereas in PPT1, PHB showed no further increase (Fig. 4a). Microscopy confirmed that, in dark-grown PPHET, PHB granules localized primarily in heterotrophic members of the community (indicated by black arrowheads in Fig. 4b), rather than in PPT1 cells (indicated with black arrows in Fig. 4b). Interestingly, not all non-cyanobacterial members contributed equally to the final PHB production. It is worth mentioning that the overall biodiversity and PHB accumulation differ consistently in Fig. 1e and Fig. 4b, indicating that while photoautotrophic and fermentative conditions both enable PHB accumulation, they shape the PPHET microbiome differently.

**Figure 4:**
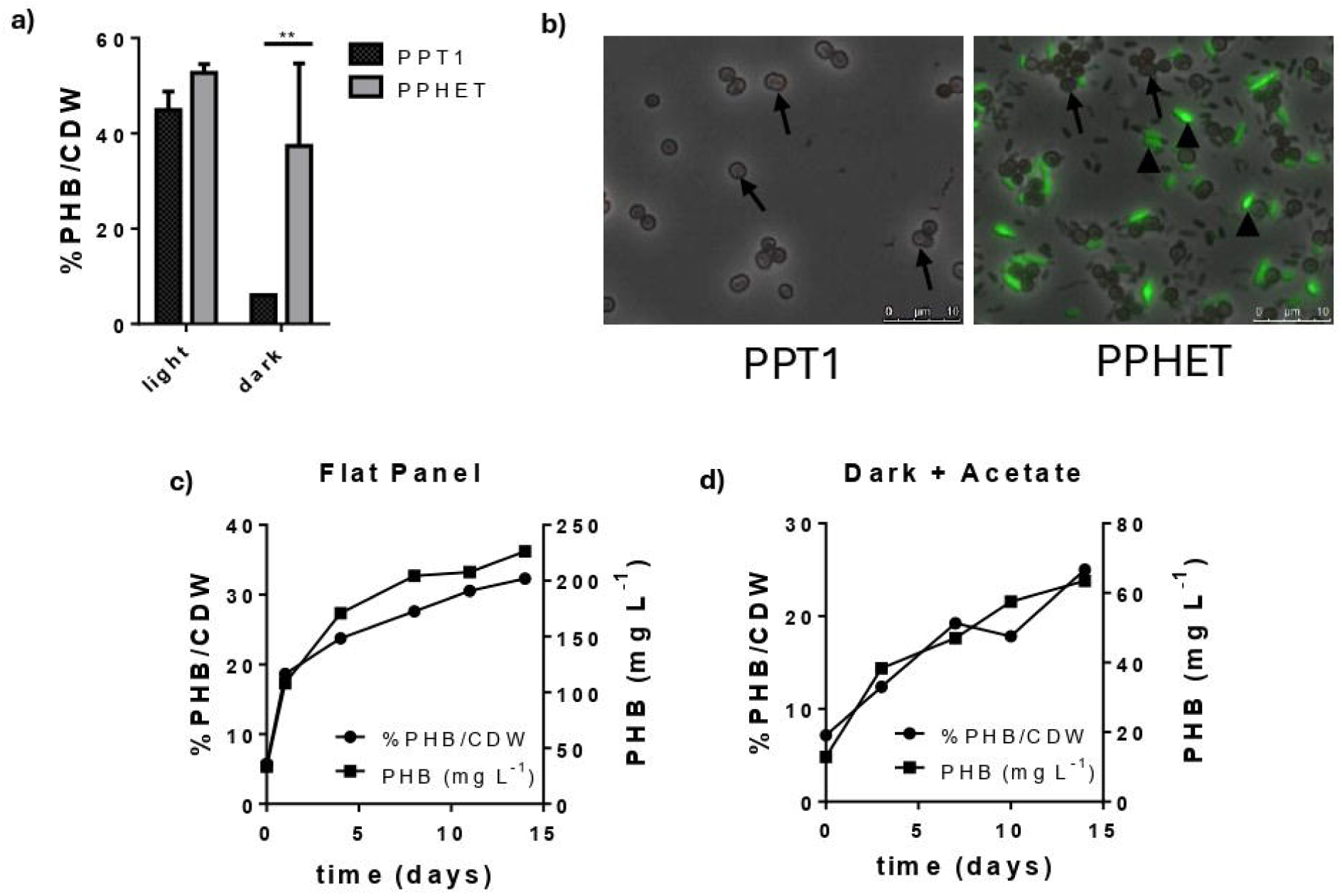
Versatility of PHB production with PPHET and upscale scenario for industrial use. **a)** PHB content in PPT1 and PPHET communities measured two weeks after 30 mM sodium acetate supplementation under light and dark conditions in small-scale cultures. Column bars represent the mean of three biological replicates ±SD. Statistical significance was determined using two-way ANOVA followed by Sidak’s multiple comparisons test. Only statistically significant differences are indicated. **b)** Fluorescence microscopy images of PPT1 and PPHET following acetate-induced PHB production in darkness. PHB granules were visualized by BODIPY staining (GFP channel) and overlaid with phase-contrast images. Arrows indicate PPT1 cells, in which no stained granules are detected in either axenic or hybrid cultures. Black arrowheads highlight non-cyanobacterial cells that contain BODIPY-stained PHB granules in PPHET. **c-d)** Time-course of PHB accumulation in the flat-panel photobioreactor system under nitrogen-limited, high-light conditions **(c)** and in the fermenter under dark conditions with acetate supplementation **(d)**. Data are presented as %PHB/CDW and total PHB titer (mg L^-1^), each dot represents a single biological replicate.

Absolute PHB titers also varied between conditions due to biomass differences. Light-incubated cultures reached OD₇₅₀ ∼3 (∼250 mg L^-1^ PHB in PPHET), whereas in darkness, cultures remained at OD₇₅₀ ∼1 (∼23 mg L^-1^ PHB) (Supporting information, Fig. S4a–b). This reflects the fact that nitrogen-depleted cyanobacterial cultures undergoing chlorosis complete a final round of cell division before entering dormancy, while in darkness, cultures did not undergo chlorosis and biomass accumulation ceased.

### Improved Scalability of Hybrid Microbiomes under Pre-Industrial Conditions

To assess whether a hybrid microbiome offers advantages compared to axenic cultures in terms of scalability, we validated our results under pre-industrial, non-sterile conditions. For this purpose, both the axenic *Synechocystis* strain PPT1 and the hybrid PPHET community were cultivated in parallel. The experiment was divided into two phases: (i) a growth phase in 10 L glass vessels, and (ii) a PHB production phase, induced either photoautotrophically via chlorosis in flat-panel reactors with high surface-to-volume ratio (to ensure sufficient light supply) or heterotrophically in dark glass reactors wrapped with aluminium foil and supplemented with acetate. Phase (ii) was inoculated directly with cultures from phase (i).

PPHET and the axenic PPT1 were grown under identical conditions, starting with BG11_0_ supplemented with 5 mM NaNO_3_ and 10 mM NaHCO_3_. Both systems initially tolerated non-sterile conditions and reached OD₇₅₀ ∼1. Upon increasing light intensity to promote further biomass accumulation and nitrate consumption, PPHET continued to grow, reaching OD₇₅₀ ∼4 (Supporting information, Fig. S5a-b). In contrast, PPT1 collapsed immediately after light increase, followed by biomass loss and complete bleaching (Supporting information, Fig. S5c). Microbial plating on contamination plates revealed atypical growth, suggesting possible contamination of the PPT1 cultures by opportunistic microbes upon death caused by environmental stress (Supporting information, Fig. S5d). These results emphasize the enhanced resilience of PPHET under scale-up conditions.

We further validated PHB production in scaled systems. Given the challenges of inducing full chlorosis in large, high-density cultures due to light limitations, PPHET was transferred into a flat-panel photobioreactor (4 L volume, 3 cm light path) and exposed to 300 µmol m⁻² s⁻¹ light in the presence of NaHCO₃. Under these conditions, PPHET reached 33% PHB/CDW (∼250 mg L^-1^) (Fig. 4c). A separate dark-scale fermentation using an 8 L, non-aerated, foil-wrapped reactor supplemented with acetate yielded 22% PHB/CDW (∼70 mg L^-1^) (Fig. 4d). Comparative analysis revealed that while PPT1 could accumulate PHB under controlled, small-scale conditions, it consistently failed under scale-up stress, unlike the robust and flexible PPHET system.

## Discussion

As a general conclusion, these results demonstrate the superior robustness and versatility of a hybrid microbiome (PPHET) compared to axenic cultures under multiple stress conditions and scaling regimes. Here, we discuss the ecological and biotechnological implications of these findings, as well as potential mechanisms underlying the enhanced performance of PPHET.

### Community composition and functional resilience of cyanobacterial consortia

Next-generation sequencing (NGS) analysis of the WCRM and PPHET communities revealed taxonomic profiles resembling those of natural cyanobacteria-dominated ecosystems. Both consortia harboured members of the order Rhodobacterales, Flavobacteriales, Rhizobiales, and Sphingomonadales, taxa commonly associated with biogeochemical cycling and symbiotic interactions (Altamira-Algarra et al., 2025b; Garcia-Pichel, 2023; Kust et al., 2025). These groups commonly co-occur with cyanobacteria in diverse environments such as in freshwaters (Kust et al., 2025), biological soil crusts (Garcia-Pichel, 2023; Nelson et al., 2021; Van Goethem et al., 2017), marine systems (Mutalipassi et al., 2021), and lichen symbioses (Aschenbrenner et al., 2016). In such communities, cyanobacteria act as primary producers, fixing carbon in nutrient-poor environments and thereby supporting the growth of other community members (Garcia-Pichel, 2023; Kust et al., 2025). These communities also include organisms capable of anoxygenic photosynthesis, such as Rhodobacterales and Sphingomonadales, suggesting that light-driven energy production is a major metabolic strategy for cyanobacteria-associated bacteria, with electron donors likely derived from organic substrates provided by cyanobacteria (Duxbury et al., 2025; Kust et al., 2025). Symbiotic relationships that facilitate the production of essential metabolites further supporting community stability and function, have also been described (Duxbury et al., 2025). In such systems, heterotrophic members consume oxygen, preventing the buildup of reactive oxygen species (ROS) and photorespiration, both of which inhibit cyanobacterial growth (Ku and Edwards, 1977). Moreover, heterotrophs are inherently more resilient to oxidative stress due to their robust ROS-scavenging systems, in contrast to cyanobacteria (Sandrini et al., 2020; Szeinbaum et al., 2021). These findings highlight how ecological principles governing the dynamics of natural microbial consortia can be leveraged to engineer communities with enhanced functionality and resilience for biotechnological applications (Giri et al., 2020).

### Robustness of the microbiome against abiotic stress factors

Fluctuations in light intensity and temperature represent major challenges for stable cultivation in industrial photobioreactors, often impairing cyanobacterial growth and productivity (Amin et al., 2024; Wahal and Viamajala, 2010). In this study, the axenic strain PPT1 exhibited sensitivity to high light, likely due to overstimulated photosynthesis leading to excessive oxygen production. This condition promotes photorespiration and the generation of reactive oxygen species (ROS), resulting in oxidative stress and reduced metabolic efficiency (Apel and Hirt, 2004; Ku and Edwards, 1977). In contrast, the synthetic community PPHET demonstrated enhanced resilience, maintaining stable growth and PHB production. This robustness is likely due to the presence of heterotrophic partners that scavenge excess oxygen, mitigating oxidative stress, a phenomenon previously reported in microalgae-associated consortia (Krohn et al., 2022).

Similarly, PPHET outperformed PPT1 under elevated temperature conditions, further supporting the protective role of the heterotrophic community in buffering environmental stress (Awasthi et al., 2014). These findings highlight the advantages of consortium-based systems for large-scale applications, where environmental fluctuations are inevitable. Overall, the enhanced resilience of PPHET aligns with the broader ecological principle that biodiversity confers stability and functional conservation under abiotic stress (Awasthi et al., 2014; Giri et al., 2020).

### Metabolic versatility in PHB production pathways – relevance for industrial applications

Nitrogen limitation is one of the most investigated conditions for PHB accumulation in cyanobacteria, typically implemented via a two-step cultivation process involving biomass generation followed by nitrogen starvation in a separate reactor (Forchhammer and Schwarz, 2019). However, effective induction of chlorosis and PHB biosynthesis requires sufficient light to activate redox-based signalling pathways (Klotz et al., 2015), posing a significant limitation for large-scale systems due to poor light penetration and high infrastructure demands (Price et al., 2022).

In this study, we demonstrate the metabolic adaptability of the PPHET synthetic microbiome, providing strategies better suited for industrial scalability. First, flat-panel photobioreactors with high surface-area illumination enabled effective chlorosis induction and PHB accumulation after nitrate depletion. This design is amenable to solar-driven systems, reducing operational complexity and costs, consistent with previous work using sunlit or open cultivation platforms (Carone et al., 2024; Price, 2022). Second, we show that PHB synthesis can also be achieved under dark conditions via acetate supplementation. This reflects fermentative PHB production by heterotrophic members of the community, providing an alternative to light-dependent, photoautotrophic accumulation. A similar dark-phase production strategy has been validated using the natural WCRM microbiome, supporting the viability of alternating light and dark cultivation modalities (Altamira-Algarra et al., 2024).

These findings underscore the value of hybrid microbiomes in enhancing operational flexibility. The ability to switch between photoautotrophic and heterotrophic production modes allows adaptation to variable cultivation environments and simplifies process control. Given the challenges of sterile, tightly regulated phototrophic cultivation (Price et al., 2022; Rueda et al., 2023; Schmelling and Bross, 2024), such microbiome-based systems present a compelling solution for scalable and economically viable bioproduction (Altamira-Algarra et al., 2025a).

### PHB quantification and productivity

In this study, we quantified PHB accumulation across different cultivation setups using multiple metrics: %PHB per cell dry weight (CDW), total PHB titer (mg L^-1^), and volumetric productivity (mg L⁻¹ d⁻¹). The primary purpose of reporting these values is not to establish absolute performance benchmarks, but to demonstrate the functional flexibility and enhanced robustness of the synthetic microbiome under diverse operational conditions.

PPHET achieved the highest PHB accumulation at small scale, reaching approximately 600 mg L^-1^, equivalent to an overall productivity of ∼35 mg L⁻¹ d⁻¹, when considering both growth and the 14-day chlorosis phase, and ∼42 mg L⁻¹ d⁻¹ when focusing solely on the chlorosis phase. For comparison, chlorosis-phase productivity in the upscaled systems was ∼18 mg L⁻¹ d⁻¹ in the flat-panel reactor (250 mg L^-1^ final yield) and ∼5 mg L⁻¹ d⁻¹ in the fermentative setup (70 mg L^-1^). These results highlight that small-scale outcomes do not directly translate to larger systems, emphasizing the need for reactor-specific performance evaluations. Nevertheless, scale-up optimization was beyond the scope of this study and represents an important avenue for future research.

## Conclusions

This study demonstrates that a natural, complex microbiome can stably host genetically engineered cyanobacteria, enabling a robust and scalable platform for phototrophic PHB production. By harnessing the ecological benefits of microbial consortia, this approach overcomes key limitations associated with axenic cultivation and offers a promising framework for industrial-scale bioprocesses. While further optimization is needed to enhance overall productivity, including strategies to increase PHB yield per biomass and improve bioreactor design on a large scale, this study shows the potential of microbiome-based approaches to advance cyanobacterial biotechnology and support the development of resilient product production platforms.

## Acknowledgments

We thank the group of Juan Garcia from the University of Catalonia for providing the WCRM, and Eva Gonzales Flo for the insights in the natural microbiome. We thank Dr. Cristoph Ratzke for the support on the initial validation of the method, and Christoph Mayer for supporting the HPLC analytics. NGS sequencing was performed at the Core Facility Genomics of the Medical Faculty, Institute for Medical Microbiology and Hygiene (MGM), University Hospital Tübingen. Data management and storage of raw data for this project were supported by the Quantitative Biology Center (QBiC), University of Tübingen, Germany. We thank Dr. Libera Lo Presti for the final proof-reading and infrastructural support through the Cluster of Excellence EXC 2124 (Controlling Microbes to Fight Infections, CMFI, grant 390838134).

## References

Altamira-Algarra, B., Garcia, J., Gonzalez-Flo, E., 2025a. Cyanobacteria microbiomes for bioplastic production: Critical review of key factors and challenges in scaling from laboratory to industry set-ups. Bioresour. Technol. 422, 132231. 10.1016/j.biortech.2025.132231

Altamira-Algarra, B., Lage, A., Meléndez, A.L., Arnau, M., Gonzalez-Flo, E., García, J., 2024. Bioplastic production by harnessing cyanobacteria-rich microbiomes for long-term synthesis. Sci. Total Environ. 954, 176136. 10.1016/j.scitotenv.2024.176136

Altamira-Algarra, B., Sun, L., San León Granado, D., Romero-Morillo, L., Vurro, L., Nogales, J., Gonzalez-Flo, E., Garcia, J., 2025b. Robust strategy for bioplastic production from cyanobacteria-enriched microbiomes: insights from gene expression and population dynamics. Chem. Eng. J. 517, 164196. 10.1016/j.cej.2025.164196

Amin, N., Jaiswal, M., Kannaujiya, V.K., 2024. Effects of temperature on morphology, physiology, and metabolic profile of diazotrophic cyanobacteria inhabiting diverse habitats. Plant Physiol. Biochem. PPB 216, 109186. 10.1016/j.plaphy.2024.109186

Apel, K., Hirt, H., 2004. REACTIVE OXYGEN SPECIES: Metabolism, Oxidative Stress, and Signal Transduction. Annu. Rev. Plant Biol. 55, 373–399. 10.1146/annurev.arplant.55.031903.141701

Aschenbrenner, I.A., Cernava, T., Berg, G., Grube, M., 2016. Understanding Microbial Multi-Species Symbioses. Front. Microbiol. 7. 10.3389/fmicb.2016.00180

Awasthi, A., Singh, M., Soni, S.K., Singh, R., Kalra, A., 2014. Biodiversity acts as insurance of productivity of bacterial communities under abiotic perturbations. ISME J. 8, 2445–2452. 10.1038/ismej.2014.91

Bolyen, E., Rideout, J.R., Dillon, M.R., Bokulich, N.A., Abnet, C.C., Al-Ghalith, G.A., Alexander, H., Alm, E.J., Arumugam, M., Asnicar, F., Bai, Y., Bisanz, J.E., Bittinger, K., Brejnrod, A., Brislawn, C.J., Brown, C.T., Callahan, B.J., Caraballo-Rodríguez, A.M., Chase, J., Cope, E.K., Da Silva, R., Diener, C., Dorrestein, P.C., Douglas, G.M., Durall, D.M., Duvallet, C., Edwardson, C.F., Ernst, M., Estaki, M., Fouquier, J., Gauglitz, J.M., Gibbons, S.M., Gibson, D.L., Gonzalez, A., Gorlick, K., Guo, J., Hillmann, B., Holmes, S., Holste, H., Huttenhower, C., Huttley, G.A., Janssen, S., Jarmusch, A.K., Jiang, L., Kaehler, B.D., Kang, K.B., Keefe, C.R., Keim, P., Kelley, S.T., Knights, D., Koester, I., Kosciolek, T., Kreps, J., Langille, M.G.I., Lee, J., Ley, R., Liu, Y.-X., Loftfield, E., Lozupone, C., Maher, M., Marotz, C., Martin, B.D., McDonald, D., McIver, L.J., Melnik, A.V., Metcalf, J.L., Morgan, S.C., Morton, J.T., Naimey, A.T., Navas-Molina, J.A., Nothias, L.F., Orchanian, S.B., Pearson, T., Peoples, S.L., Petras, D., Preuss, M.L., Pruesse, E., Rasmussen, L.B., Rivers, A., Robeson, M.S., Rosenthal, P., Segata, N., Shaffer, M., Shiffer, A., Sinha, R., Song, S.J., Spear, J.R., Swafford, A.D., Thompson, L.R., Torres, P.J., Trinh, P., Tripathi, A., Turnbaugh, P.J., Ul-Hasan, S., Van Der Hooft, J.J.J., Vargas, F., Vázquez-Baeza, Y., Vogtmann, E., Von Hippel, M., Walters, W., Wan, Y., Wang, M., Warren, J., Weber, K.C., Williamson, C.H.D., Willis, A.D., Xu, Z.Z., Zaneveld, J.R., Zhang, Y., Zhu, Q., Knight, R., Caporaso, J.G., 2019. Reproducible, interactive, scalable and extensible microbiome data science using QIIME 2. Nat. Biotechnol. 37, 852–857. 10.1038/s41587-019-0209-9

Callahan, B.J., McMurdie, P.J., Rosen, M.J., Han, A.W., Johnson, A.J.A., Holmes, S.P., 2016. DADA2: High-resolution sample inference from Illumina amplicon data. Nat. Methods 13, 581–583. 10.1038/nmeth.3869

Caporaso, J.G., Lauber, C.L., Walters, W.A., Berg-Lyons, D., Lozupone, C.A., Turnbaugh, P.J., Fierer, N., Knight, R., 2011. Global patterns of 16S rRNA diversity at a depth of millions of sequences per sample. Proc. Natl. Acad. Sci. U. S. A. 108 Suppl 1, 4516–4522. 10.1073/pnas.1000080107

Carone, M., Frungieri, G., Costamagna, L., Zanetti, M., Vanni, M., Riggio, V.A., 2024. Advanced Design and Characterization of a Flat Panel Photobioreactor Equipped with a Customizable Light-Emitting Diode Lighting System. 10.1021/acssuschemeng.3c05176

Da Veiga Leprevost, F., Grüning, B.A., Alves Aflitos, S., Röst, H.L., Uszkoreit, J., Barsnes, H., Vaudel, M., Moreno, P., Gatto, L., Weber, J., Bai, M., Jimenez, R.C., Sachsenberg, T., Pfeuffer, J., Vera Alvarez, R., Griss, J., Nesvizhskii, A.I., Perez-Riverol, Y., 2017. BioContainers: an open-source and community-driven framework for software standardization. Bioinformatics 33, 2580–2582. 10.1093/bioinformatics/btx192

Doello, S., Burkhardt, M., Forchhammer, K., 2021. The essential role of sodium bioenergetics and ATP homeostasis in the developmental transitions of a cyanobacterium. Curr. Biol. 31, 1606–1615.e2. 10.1016/j.cub.2021.01.065

Duxbury, S.J.N., Raguideau, S., Cremin, K., Richards, L., Medvecky, M., Rosko, J., Coates, M., Randall, K., Chen, J., Quince, C., Soyer, O.S., 2025. Niche formation and metabolic interactions contribute to stable diversity in a spatially structured cyanobacterial community. ISME J. 19, wraf126. 10.1093/ismejo/wraf126

Ewels, P., Peltzer, A., Fillinger, S., Patel, H., Alneberg, J., Wilm, A., Garcia, M.U., Di Tommaso, P., Nahnsen, S., 2022. The nf-core framework for community-curated bioinformatics pipelines. 10.5281/zenodo.7153103

Ewels, P.A., Peltzer, A., Fillinger, S., Patel, H., Alneberg, J., Wilm, A., Garcia, M.U., Di Tommaso, P., Nahnsen, S., 2020. The nf-core framework for community-curated bioinformatics pipelines. Nat. Biotechnol. 38, 276–278. 10.1038/s41587-020-0439-x

Fink, P., Menzel, C., Kwon, J.-H., Forchhammer, K., 2025. A novel recombinant PHB production platform in filamentous cyanobacteria avoiding nitrogen starvation while preserving cell viability. Microb. Cell Factories 24, 43. 10.1186/s12934-025-02650-y

Forchhammer, K., Schwarz, R., 2019. Nitrogen chlorosis in unicellular cyanobacteria – a developmental program for surviving nitrogen deprivation. Environ. Microbiol. 21, 1173–1184. 10.1111/1462-2920.14447

Gao, H., Manishimwe, C., Yang, L., Wang, H., Jiang, Y., Jiang, W., Zhang, W., Xin, F., Jiang, M., 2022. Applications of synthetic light-driven microbial consortia for biochemicals production. Bioresour. Technol. 351, 126954. 10.1016/j.biortech.2022.126954

Garcia-Pichel, F., 2023. The Microbiology of Biological Soil Crusts. Annu. Rev. Microbiol. 77, 149–171. 10.1146/annurev-micro-032521-015202

Gautam, S., Gautam, A., Pawaday, J., Kanzariya, R.K., Yao, Z., 2024. Current Status and Challenges in the Commercial Production of Polyhydroxyalkanoate-Based Bioplastic: A Review. Processes 12, 1720. 10.3390/pr12081720

Giri, S., Shitut, S., Kost, C., 2020. Harnessing ecological and evolutionary principles to guide the design of microbial production consortia. Curr. Opin. Biotechnol. 62, 228–238. 10.1016/j.copbio.2019.12.012

Grüning, B., Dale, R., Sjödin, A., Chapman, B.A., Rowe, J., Tomkins-Tinch, C.H., Valieris, R., Köster, J., Bioconda Team, 2018. Bioconda: sustainable and comprehensive software distribution for the life sciences. Nat. Methods 15, 475–476. 10.1038/s41592-018-0046-7

Gundlapalli, M., Ganesan, S., 2025. Polyhydroxyalkanoates (PHAs): Key Challenges in production and sustainable strategies for cost reduction within a circular economy framework. Results Eng. 26, 105345. 10.1016/j.rineng.2025.105345

Huang, Q., Jiang, F., Wang, L., Yang, C., 2017. Design of Photobioreactors for Mass Cultivation of Photosynthetic Organisms. Engineering 3, 318–329. 10.1016/J.ENG.2017.03.020

Jousset, A., Schulz, W., Scheu, S., Eisenhauer, N., 2011. Intraspecific genotypic richness and relatedness predict the invasibility of microbial communities. ISME J. 5, 1108–1114. 10.1038/ismej.2011.9

Klotz, A., Reinhold, E., Doello, S., Forchhammer, K., 2015. Nitrogen Starvation Acclimation in Synechococcus elongatus: Redox-Control and the Role of Nitrate Reduction as an Electron Sink. Life 5, 888–904. 10.3390/life5010888

Koch, M., Bruckmoser, J., Scholl, J., Hauf, W., Rieger, B., Forchhammer, K., 2020. Maximizing PHB content in Synechocystis sp. PCC 6803: a new metabolic engineering strategy based on the regulator PirC. Microb. Cell Factories 19, 231. 10.1186/s12934-020-01491-1

Krohn, I., Menanteau-Ledouble, S., Hageskal, G., Astafyeva, Y., Jouannais, P., Nielsen, J.L., Pizzol, M., Wentzel, A., Streit, W.R., 2022. Health benefits of microalgae and their microbiomes. Microb. Biotechnol. 15, 1966–1983. 10.1111/1751-7915.14082

Ku, S.-B., Edwards, G.E., 1977. Oxygen Inhibition of Photosynthesis: II. Kinetic Characteristics as Affected by Temperature 1. Plant Physiol. 59, 991–999. 10.1104/pp.59.5.991

Kust, A., Zorz, J., Paniker, C.C., Bouma-Gregson, K., Krishnappa, N., Liu, W., Banfield, J.F., Diamond, S., 2025. Model cyanobacterial consortia reveal a consistent core microbiome independent of inoculation source or cyanobacterial host species. ISME J. 19, wraf142. 10.1093/ismejo/wraf142

McAdam, B., Brennan Fournet, M., McDonald, P., Mojicevic, M., 2020. Production of Polyhydroxybutyrate (PHB) and Factors Impacting Its Chemical and Mechanical Characteristics. Polymers 12, 2908. 10.3390/polym12122908

Mutalipassi, M., Riccio, G., Mazzella, V., Galasso, C., Somma, E., Chiarore, A., de Pascale, D., Zupo, V., 2021. Symbioses of Cyanobacteria in Marine Environments: Ecological Insights and Biotechnological Perspectives. Mar. Drugs 19, 227. 10.3390/md19040227

Nelson, C., Giraldo-Silva, A., Garcia-Pichel, F., 2021. A symbiotic nutrient exchange within the cyanosphere microbiome of the biocrust cyanobacterium, *Microcoleus vaginatus*. ISME J. 15, 282–292. 10.1038/s41396-020-00781-1

Orthwein, T., Scholl, J., Spät, P., Lucius, S., Koch, M., Macek, B., Hagemann, M., Forchhammer, K., 2021. The novel PII-interactor PirC identifies phosphoglycerate mutase as key control point of carbon storage metabolism in cyanobacteria. Proc. Natl. Acad. Sci. U. S. A. 118, e2019988118. 10.1073/pnas.2019988118

Price, S., 2022. Developing Next Generation Algae Bioplastic Technology (Ph.D.). PQDT - Glob. University of Technology Sydney (Australia), Australia.

Price, S., Kuzhiumparambil, U., Pernice, M., Ralph, P., 2022. Techno-economic analysis of cyanobacterial PHB bioplastic production. J. Environ. Chem. Eng. 10, 107502. 10.1016/j.jece.2022.107502

Quast, C., Pruesse, E., Yilmaz, P., Gerken, J., Schweer, T., Yarza, P., Peplies, J., Glöckner, F.O., 2012. The SILVA ribosomal RNA gene database project: improved data processing and web-based tools. Nucleic Acids Res. 41, D590–D596. 10.1093/nar/gks1219

Rippka, R., Deruelles, J., Waterbury, J.B., Herdman, M., Stanier, R.Y., 1979. Generic Assignments, Strain Histories and Properties of Pure Cultures of Cyanobacteria. Microbiology 111, 1–61. 10.1099/00221287-111-1-1

Rueda, E., Senatore, V., Zarra, T., Naddeo, V., García, J., Garfí, M., 2023. Life cycle assessment and economic analysis of bioplastics production from cyanobacteria. Sustain. Mater. Technol. 35, e00579. 10.1016/j.susmat.2023.e00579

Salazar, V.W., Tschoeke, D.A., Swings, J., Cosenza, C.A., Mattoso, M., Thompson, C.C., Thompson, F.L., 2020. A new genomic taxonomy system for the Synechococcus collective. Environ. Microbiol. 22, 4557–4570. 10.1111/1462-2920.15173

Sandrini, G., Piel, T., Xu, T., White, E., Qin, H., Slot, P.C., Huisman, J., Visser, P.M., 2020. Sensitivity to hydrogen peroxide of the bloom-forming cyanobacterium *Microcystis* PCC 7806 depends on nutrient availability. Harmful Algae 99, 101916. 10.1016/j.hal.2020.101916

Schmelling, N.M., Bross, M., 2024. What is holding back cyanobacterial research and applications? A survey of the cyanobacterial research community. Nat. Commun. 15, 6758. 10.1038/s41467-024-50828-6

Straub, D., Blackwell, N., Langarica-Fuentes, A., Peltzer, A., Nahnsen, S., Kleindienst, S., 2020. Interpretations of Environmental Microbial Community Studies Are Biased by the Selected 16S rRNA (Gene) Amplicon Sequencing Pipeline. Front. Microbiol. 11, 550420. 10.3389/fmicb.2020.550420

Szeinbaum, N., Toporek, Y.J., Reinhard, C.T., Glass, J.B., 2021. Microbial helpers allow cyanobacteria to thrive in ferruginous waters. Geobiology 19, 510–520. 10.1111/gbi.12443

Van Goethem, M.W., Makhalanyane, T.P., Cowan, D.A., Valverde, A., 2017. Cyanobacteria and Alphaproteobacteria May Facilitate Cooperative Interactions in Niche Communities. Front. Microbiol. 8, 2099. 10.3389/fmicb.2017.02099

Wahal, S., Viamajala, S., 2010. Maximizing algal growth in batch reactors using sequential change in light intensity. Appl. Biochem. Biotechnol. 161, 511–522. 10.1007/s12010-009-8891-6

