## Supplementary materials for "Cultivation in a Natural Microbial Community Enhances the Industrial Performance of a Genetically Engineered Cyanobacterium for Bioplastic Production"

### Supporting information

**Table 1.** List of the cultures used in this study.

| NAME | GENOTYPE | REFERENCE |
|---|---|---|
| PPT1 | *Synechocystis* sp. PCC 6803, KanR, GenR | Koch *et al.*, 2020 |
| GFP | *Synechocystis* sp. PCC 6803, SpecR | Orthwein *et al.*, 2021 |
| WRCM | Wild Cyanobacteria Rich Microbiome | Altamira-Algarra *et al.*, 2024 |
| PPHET | PPT1 with non-cyanobacterial part of WCRM | This study |

**Table 2.** List of the microorganisms grouped in the group “Others” in Fig. 2b, with relative abundancy lower than 0,05% at the time of sampling.

| PHYLUM | CLASS | ORDER | NOTES |
|---|---|---|---|
| Actinobacteriota | Actinobacteria | Micrococcales | |
| Myxococcota | Myxococcia | Myxococcales | |
| Proteobacteria | Gammaproteobacteria | Enterobacterales | |
| Proteobacteria | Gammaproteobacteria | Burkholderiales | |
| Cyanobacteria | Cyanobacteriia | Phormidesmiales | present only in WCRM |
| Acidobacteriota | Acidobacteriae | Bryobacterales | |
| Cyanobacteria | Cyanobacteriia | Leptolyngbyales | present only in WCRM |
| Verrucomicrobiota | Verrucomicrobiae | Chthoniobacterales | present only in WCRM |

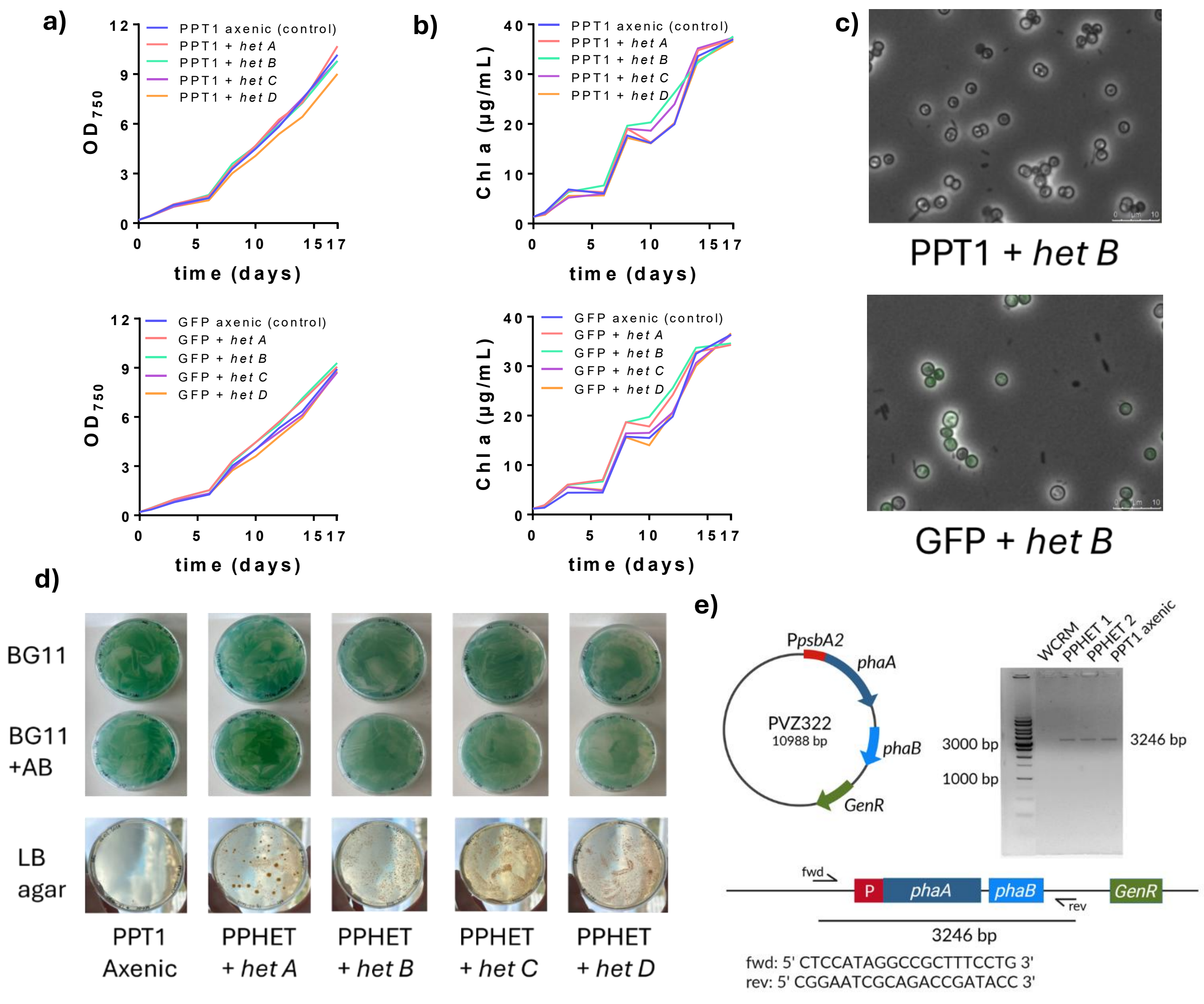


**Figure S1: Hybrid microbiome establishment and stability. a-b)** Growth ($OD_{750}$) and chlorophyll *a* accumulation were monitored over time to specifically track the phototrophic component (cyanobacteria) within the synthetic communities; **b)** Microscopic visualization of the new microbiomes. Images show overlays of phase-contrast and GFP fluorescence channels to enable simultaneous observation of overall cellular morphology and on the right, the distribution of GFP-expressing cyanobacteria within the community; **c)** To confirm that PPT1 remained the dominant phototroph within the community, cultures were plated on BG11 agar both with and without the addition of antibiotics to which PPT1 is resistant (Kanamycin, specifically). Growth under selective conditions confirmed stable integration of the engineered cyanobacterium into the microbiome. Community structure was qualitatively assessed by plating on nutrient-rich medium (LB) to evaluate heterotrophic fraction; **d)** To assess long-term community stability, cultures were propagated through multiple 14-day cycles, each followed by a 1:10 dilution into fresh BG11 medium. **e)** To verify the continued presence of the engineered strain and exclude residual presence and overgrowth by wild-type *Synechocystis*, colony PCR targeting the PPT1 plasmid was performed at multiple time points. The plasmid is reported with the Promotor *psbA2* (P*psbA2*) from *Synechocystis* and *phaAB* gene cassette from *Cupriavidus necator* H16, which were assembled in the PVZ322 vector (Koch *et al,* 2020). The linearized plasmid is represented at the bottom of the picture, and semi-arrows correspond to primers annealing to it (primer sequences reported). No amplification was observed in the wild-type community (WRCM), while distinct bands corresponding to *phaA* and *phaB* were detected in the PPHET community after initial inoculation (PPHET 1) and after the second propagation cycle (PPHET 2). A positive control from the axenic PPT1 culture is also shown. This result confirms successful integration of the engineered strain PPT1 within the hybrid community.

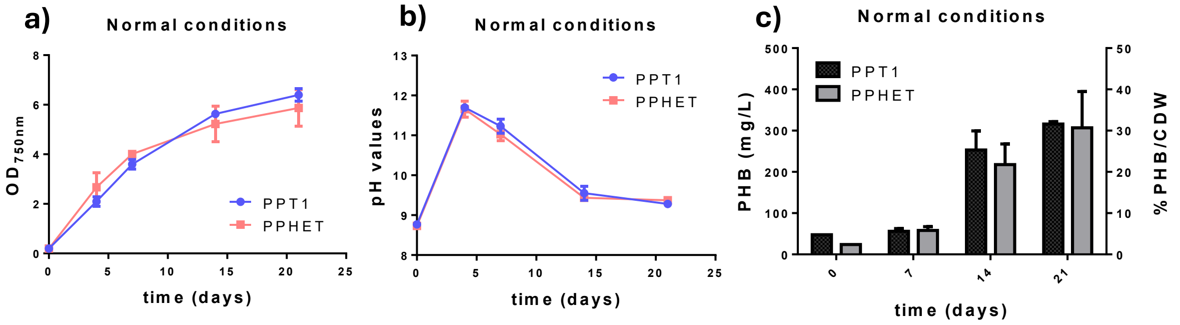


**Figure S2: PPT1 and PPHET in normal conditions as a control to the stress test do not show big differences.** **a)** Growth curve measured at $OD_{750nm}$. **b)** pH monitoring. **c)** PHB production of PPT1 and PPHET over time.

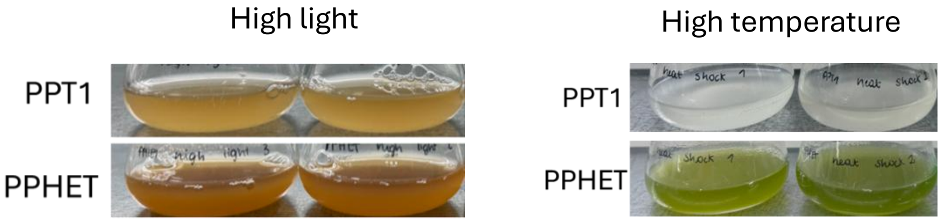


**Figure S3: PPT1 shows weaker or bleached phenotype after abiotic stresses.** Pictures of the cultures PPT1 and PPHET (duplicates) taken at the end of each experiment under high light (left) and under “heat shock” (right).

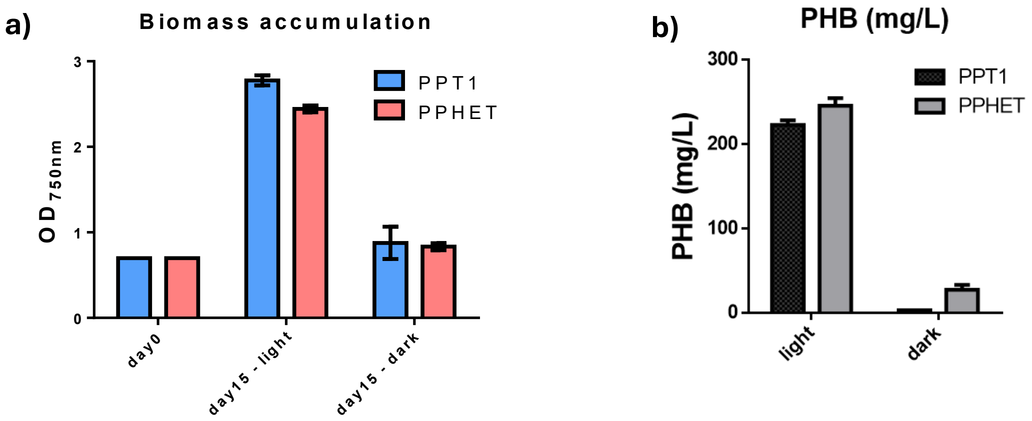


**Figure S4: The biomass accumulation influences the final PHB titer.** Cultures were analysed after 14 days upon addition of acetate, both under light and dark conditions. **a)** Biomass accumulation in $OD_{750}$. **b)** PHB production in terms of mg/L.

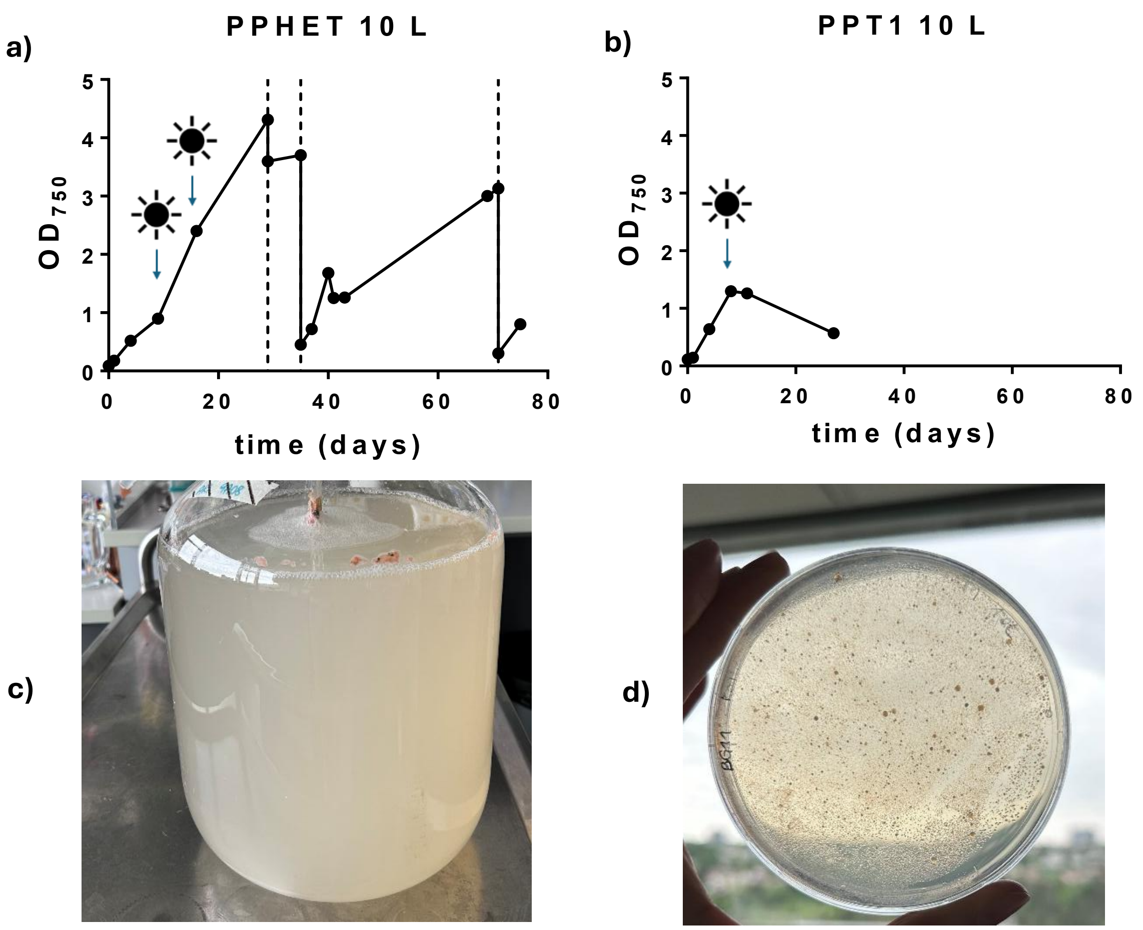


**Figure S5: the abiotic perturbances in large volumes cause the collapse of the axenic PPT1 culture. a)** Long-term cultivation of the PPHET consortium in a 10 L photobioreactor over ~80 days. Optical density ($OD_{750}$) was monitored to assess growth dynamics. Sun symbols indicate steps of increased light intensity. Vertical dotted lines in the graphs indicate the time points at which the cultures completely consumed the initial nitrate supply, marking nitrate depletion events. These were followed by partial harvesting of the cultures and replenishment with fresh $BG11_0$ medium supplemented with 1 to 4 mM $NaNO_3$. **b)** Growth profile of axenic PPT1 strain in a 10 L photobioreactor. A sharp decline in biomass was observed following increased light intensity, indicating light-induced culture collapse. **c)** Collapsed PPT1 culture in the 10 L vessel. **d)** Growth of the collapsed culture on solid media to check for contamination.

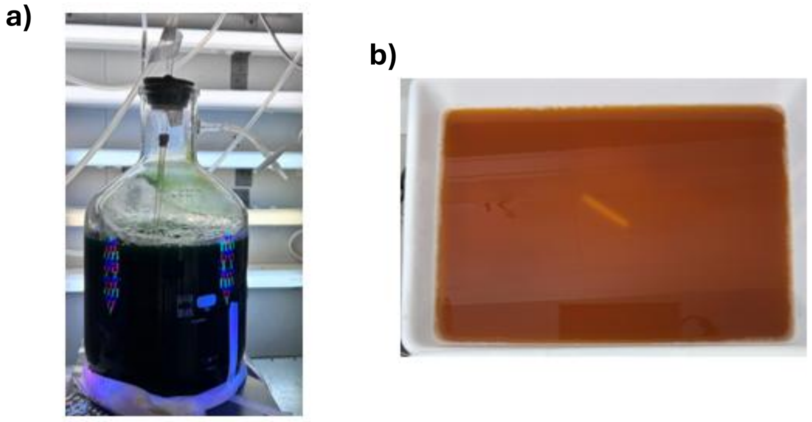


**Figure S6: Successful scaling up attempts with PPHET for growth and PHB production. a)** PPHET in the 10 L flask for normal growth. **b)** Flat Panel reactor with a 4 L volume at the end of the production process.
